# Multicolor fluorescent imaging by space-constrained computational hyperspectral imaging

**DOI:** 10.1101/458869

**Authors:** Yina Wang, Bin Yang, Siyu Feng, Veronica Pessino, Bo Huang

## Abstract

Hyperspectral imaging is a powerful technique to simultaneously study multiple fluorophore labels with overlapping emissions. Here we present a computational hyperspectral imaging method, which uses the sample spatial fluorescence information as a reconstruction constraint. Our method addresses both the under-sampling issue of compressive hyperspectral imaging and the low throughput issue of scanning hyperspectral imaging. With simulated and experimental data, we have demonstrated the superior reconstruction precision of our method in two and three-color imaging. We have experimentally validated this method in differentiating cellular structures labeled with two red-colored fluorescent proteins, tdTomato and mCherry, which have highly overlapping emission spectra. Our method has the advantage of totally free wavelength choice and can also be combined with conventional filter-based sequential multi-color imaging to further expand the choices of probes.

## 1. Introduction

In fluorescence microscopy, it is desirable to selectively label sample structures with differently colored fluorophores to study interactions [1]. Commonly, multiple colors (namely channels) are either imaged sequentially by using different filter sets or simultaneously by splitting signals onto different regions of the camera or onto several cameras. Both approaches rely on filters and are therefore ultimately limited by the spectral overlap of fluorophores, which makes it difficult in practice to distinguish more than four colors within the visible spectrum without having substantial crosstalk among channels. Sequential imaging of more than four targets is particularly challenging for live samples.

As an alternative, hyperspectral imaging is a powerful tool to simultaneously study multiple labels in biological samples at the subcellular, cellular and tissue level [1-3]. Traditionally, in hyperspectral imaging, a three-dimensional (3D) data cube (2D spatial, 1D spectral) is generated via spatial or wavelength scanning (hereafter called scanning hyperspectral imaging), where a diffraction grating or prism is used to acquire the full spectrum at each spatial point of the image [4]. With the spectral information, hyperspectral imaging approaches are capable of unambiguously identifying fluorophores with overlapping spectra and permitting high levels of signal multiplexing [5]. However, a major drawback of scanning hyperspectral imaging is the relatively long acquisition time. To solve this problem, snapshot hyperspectral imaging methods that based on tomographic multiplexing [6, 7] or compressive hyperspectral imaging [8-10] were developed. In the later, the entire data cube was either coded and captured in a single 2D camera integration with different wavelength signal dispersed to different spatial locations [8, 9] or coded by a series of Hadamard patterns to generate single pixel signals[10], and then the whole data cube was reconstructed using compressed sensing theory. Unfortunately, these methods reportedly are difficult to be applied to high resolution imaging [4] and only multicolor fluorescent beads images have been demonstrated [9, 10].

In this paper, we demonstrate a computational hyperspectral imaging method, which aims to address both the under-sampling issue of compressive hyperspectral imaging and the low throughput issue of scanning hyperspectral imaging. To do so, we employed two strategies: (1) using a dual-view imaging system to generate both an undispersed spatial image capturing the superposition of multiple labels and a spectral image where signals were dispersed by a wedge prism to different spatial locations according to wavelength (Figs. 1a and b); (2) using Digital Micromirror Device (DMD) [8, 9] to generate multiple randomly coded illumination pattern on the sample. In our approach, the undispersed spatial image acted as a spatial constraint in data reconstruction to guarantee correct reconstruction and enhance accuracy. This strategy is similar to what was used in compressed ultrafast photography [11, 12]. With both simulation and experimental data, we demonstrated that for two-color imaging, strategy (1) alone ensures a high reconstruction accuracy in resolving labels with emission peaks separated by ~20 nm. Using simulation, we also show that for three-color imaging, the use of five randomly coded illumination patterns is sufficient for an accurate reconstruction, which is still much faster than scanning hyperspectral imaging.

**Fig. 1.**
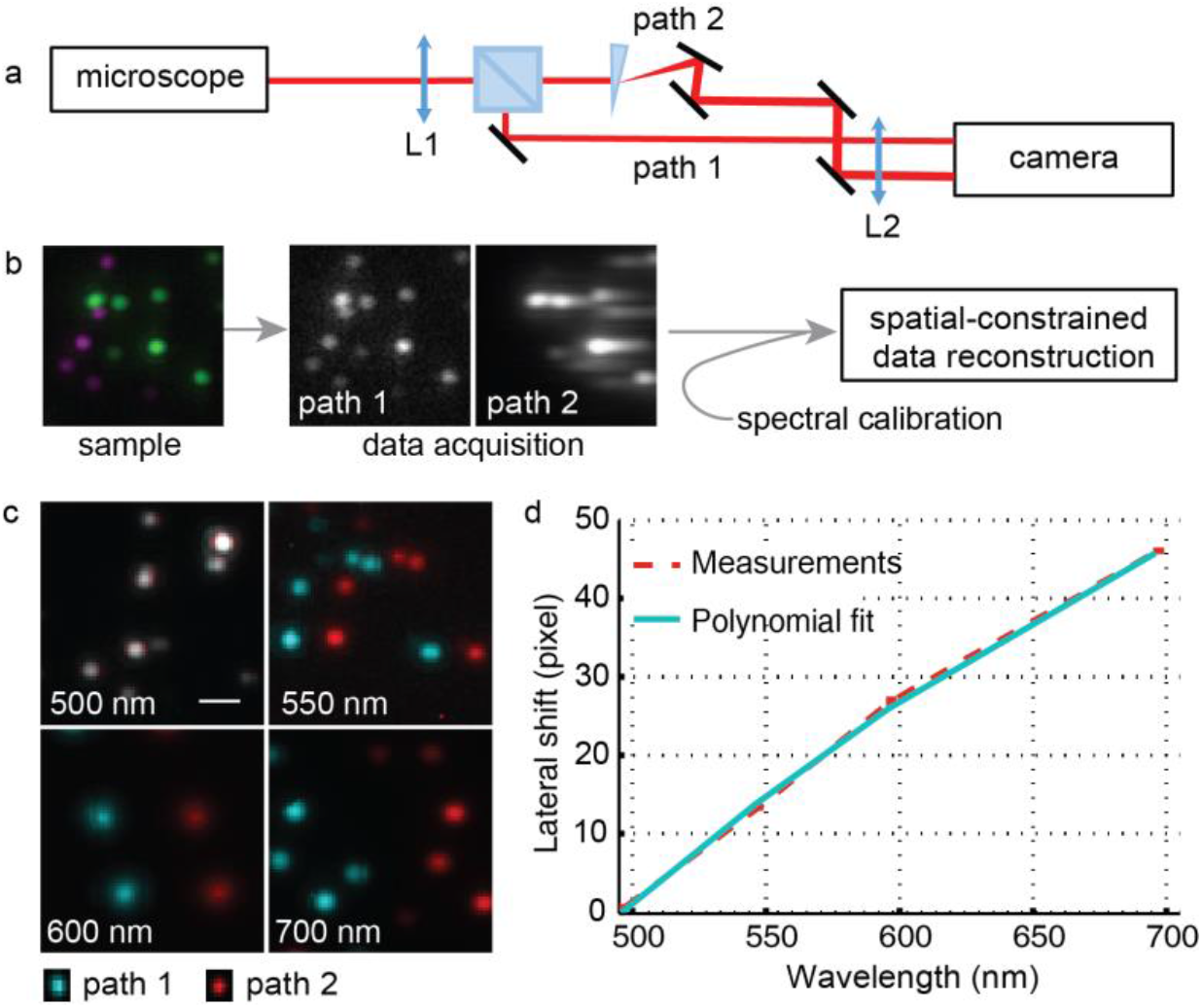
Space-constrained computational hyperspectral imaging detection system. (a) Schematic of the setup. The emitting light from the sample is split evenly by a beam splitter to generate an undispersed spatial image (Path 1) and a dispersed spectral image (Path 2) respectively on the left and right parts of the same camera. L, lens. (b) Illustration of overall space-constrained computational hyperspectral imaging approach using green and red fluorescent beads. (c) Registered spatial (blue) and spectral (red) images of fluorescent beads at different wavelength. Scale bar: 1 μm. (d) Measured lateral spatial shift and its corresponding polynomial fitting. The measurement is averaged from 5 independent experiments, with a negligible standard deviation.

## 2. Method

### 2.1 Space-constrained computational hyperspectral imaging

We aim to image a three-dimensional sample, ***x*** ={ *x*(*i*, *j*, *k*); *i*, *j* = 1 … *N*, *k* = 1 … *M* }, a distribution of fluorescent probes with *x*(*i*, *j*, *k*) represents the intensity of the *k*_*th*_ kind of probes at spatial pixel localization (*i*, *j*), *N* is the number of pixels along one spatial dimension (for simplicity, we assumed the imaged filed-of-view to be square) and *M* is the number of probes used to label the sample.

To achieve high imaging quality of the sample, we simultaneously measure its spatial and spectral information. Here, we employed a dual-view detection scheme (Fig. 1a), which is similar to what used in spectral-resolved super-resolution localization microscopy and spectral single molecule tracking [13-16]. In the detection path, the emitted fluorescence was split by a 50:50 beam splitter cube (BS013, Thorlabs). The reflected fluorescence was directed to the camera by a mirror (Path 1, referred to as the spatial path) to generate a spatial image ***y***_1_. The transmitted fluorescence passed through a dispersive wedge prism (SM1W189, Thorlabs), which shifts blue to red emission from left to right along lateral direction, and two pairs of mirrors (Path 2, referred to as spectral path) to produce a spectral image ***y***_2_. With spectral calibration results (Section 2.2), ***x*** can be reconstructed through solving an optimization problem in which ***y***_1_ acts as a spatial constraint in restoring the spectral information contained in ***y***_2_ (Fig. 1b).

Here, we first describe the forward imaging formation model of our system. In the spatial path, ***y***_1_ is the superposition integral of all probe signals at each spatial location, which is expressed as 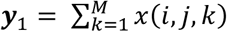. ***y***_1_ can be described in matrix form:

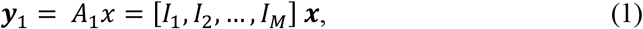

in which ***y***_1_ and ***x*** are the vectorized form, *I* is *N* × *N* identical matrix. On the other hand, in the spectral path, because we model the third dimension of ***x*** as the choice of probes instead of wavelength, the formation of ***y***_2_ can be described as a convolution problem, in which the emission spectrum of each probe and the dispersion by the prism determine the corresponding convolution kernel function. Assuming *h*(*k*) is the dispersed convolution kernel function for the *k_th_* channel (obtained through spectral calibration), then ***y***_2_ is the summation of convolved signals: 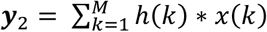. This convolution can also be expressed in matrix format with each column of the measurement matrix, col_i_*A*_2_ (*i* = 1 … *N* × *N* × *M*), corresponding to the convolved image assuming a single probe is placed at *i* position of vectorized ***x***:

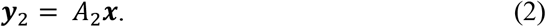

Combining Eq. (1) and (2), we have:

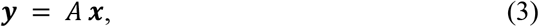

with ***y*** = [***y***_1_; ***y***_2_] and ***A*** =[***A***_1_; ***A***_2_].

To reconstruct ***x*** under the forward imaging model (Fig. 1b), two scenarios need to be considered. First, when the sample contains only two colors, Eq. (3) is a determined equation. Estimating ***x*** is a deconvolution problem. However, because of the ill-conditioned nature of deconvolution problem, general inverse matrix method cannot be used. Therefore, we need to solve the inversion problem with regularization terms to stabilize the solution:

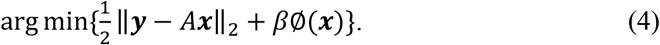

In our implementation, we adopted the two-step iterative shrinkage/thresholding (TwIST) [17] algorithm, and chose ∅(*x*) in the form of summation of 2D total variation (TV) of each channel:

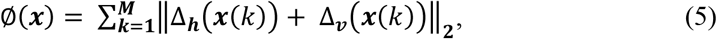

Δ*_h_* and Δ*_v_* are the horizontal and vertical first-order local difference operators. In Eq. (4), the first term, ‖***y*** − *A****x*** ‖_2_, is the fidelity term which encourages the actual measurement ***y*** to closely match the estimated measurement *A****x***. The second term, *β*∅(***x***), is the regularization term, which encourages ***x*** to be piecewise constant (that is, sparse in the gradient domain). The weighting of these two terms is empirically adjusted by the regularization parameter, *β*, to lead to results that are consistent with the physical reality. In both our simulation and experiments, we found a *β* value between 1 to 5 gave good reconstruction results. In practice, the problem is also subjective to non-negative constraint to meet physical reality requirement.

In a second scenario, when the sample is multi-color labeled (*M* > 2) and only one snapshot is taken, the number of variables in *x* becomes larger than the number of measurements. In this case, solving Eq. (3) becomes a compressive sensing problem [18]. Due to the overall similarity of emission spectrum of fluorescence proteins, the measurement matrix *A* would generally lack the incoherence required for successful compressive sensing reconstruction. This problem can be solved by acquiring multiple measurements of the sample using random illumination patterns generated by DMD. Under this strategy, the forward imaging model becomes:

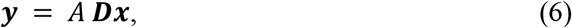

in which ***D*** is a binary matrix describing the random illumination pattern. **y** becomes the cascade of the multiple measurements, **A** is also a cascade of the original measurement matrix with certain columns setting to zero according the corresponding illumination pattern. This strategy guarantees a successful reconstruction in two ways: (1) the incoherence of the measurement matrix *A* increases because of the randomized illumination; (2) with more measurements, Eq. (3) could become an over-determined problem. Thus, ***x*** can be obtained by solving the same problem of Eq. (4) with ***AD*** being the new measurement matrix. We will illustrate the setup in Section 3.3.

### 2.2 Spectral calibration

Calibration between the spectral and spatial paths was performed using fluorescent beads in different colors (Yellow-green, orange, red, and dark red, F10720, Life Technologies). Each kind of beads was deposited to the coverslip at low density. Upon imaging, individual beads appeared as well-resolved diffraction-limited spots (Fig. 1b). Narrow bandpass filters (FB500-10, FB550-10, FB600-10, FB700-10, Thorlabs) were placed between beam splitter and the wedge prism to determine the spectral positions of the corresponding wavelength in the spectral image relative to the image position in the spatial image. Then the paired images were registered (described in 2.3 below, Fig. 1b) to estimate the lateral spatial shift of different wavelength using cross-correlation. The measured lateral shift curve can further be fitted with a two-order polynomial function to generation the prism dispersion function (Fig. 1c). This calibration only needs to be performed once for all experiments performed on the setup.

In order to generate the measurement matrix A, we need the prior knowledge of the emission spectra of probes to firstly generate the convolution kernel functions in A. For this purpose, there are usually two methods. One method is using the probe emission spectrum from literatures and calculate its corresponding dispersed convolution kernel function using the calibrated prism dispersion function. The second approach is to calibrate convolution kernel functions directly using single color images. In this paper, we used the second method because it results in higher reconstruction accuracy with systematic error included. In the single-color imaging scenario, we can also use forward imaging model to obtain the convolution kernel function. In this case, ***y*** is the spectral measurement from single-color labeled sample, ***x*** is the one-dimensional convolution kernel function, *A* is a matrix with each of its column representing the corresponding lateral shifted spatial measurement. Because the size of ***x*** is much smaller than ***y***, the overdetermined problem ***y*** = ***Ax*** can be solved by the generalized inverse matrix method ***x*** = (***A***^*T*^***A***) ^−1^ ***A***^**T**^***y***.

### 2.3 Dual-view image registration

Image registration was performed before calibration and reconstruction to avoid dual-view mismatch and the effect of potential optical aberrations. We used control point-based image registration method. First, yellow-green fluorescent beads were imaged at 500 nm. Well-separated spots, which acted as control-points, were localized using home-built C++ software [19]. Then the localized beads positions were manually checked to identify four pairs of matched spots from the four corners of the spatial and spectral images. The positions of the four pairs of spots were used to calculate the initial registration parameters from the spectral channel to the spatial channel. After applying the initial transformation, the program automatically identified all matched spots and determined a three-order polynomial coordinate transform function between the two images with a least-square fitting to the coordinates of these matched spots. The polynomial function is: *x_c_* = *A*_0_ + *A*_1_*x* + *A*_2_*y* + *A*_3_*x*^2^ + *A*_4_*xy* + *A*_5_*y*^2^ + *A*_6_*x*^3^ + *A*_7_*xy*^2^ + *A*_8_*x*^2^*y* + *A*_9_*y*^3^; *y_c_* = *B*_0_ + *B*_1_*y* + *B*_2_*x* + *B*_3_*y*^2^ + *B*_4_*xy* + *B*_5_*x*^2^ + *B*_6_*y*^3^ + *B*_7_*x*^2^*y* + *B*_8_*xy*^2^ + *B*_9_*x*^3^, where *x_c_* and *y_c_* are the transformed spectral coordinates, *x* and *y* are the initial spectral coordinates, and *A_n_* and *B_n_* are coefficients obtained by least-square fitting. The described procedure was repeated on five independently measured datasets to generate an averaged registration function. Finally, backward registration [20] from spatial channel to spectral channel was performed to eliminate registration artifact.

### 2.4 Simulation

Simulated datasets were used to quantitatively evaluate the performance of the image reconstruction method. The data sets were based on experimental images of various organelles acquired previously (ground truth images) and generated by the following steps: (1) for each image, we assume it was generated by a fluorophore with known emission spectrum (downloaded from Fluorescence SpectraViewer (ThermoFisher)); (2) These emission spectra and prism dispersion function were used firstly to generate convolution kernel functions for each choosen probe, and then to build the measurement matrix ***A***; (3) The spatial and spectral images were generated by using Eq. (1) and (2); (4) Poisson noise was added to both images to generate the synthesized image data sets. To generate datasets with various signal-to-noise ratio (SNR) and signal-to-background ratio (SBR), ground truth images were normalized to its corresponding mean intensity value and then multiplied with desired intensity level. Background was homogeneously added to the images according to desired SBR.

### 2.5 Sample preparation and imaging

BSC-1 cells (African green monkey kidney cells, from UCSF Cell Culture Facility) were maintained in Dulbecco’s modified Eagle medium (DMEM) with high glucose (UCSF Cell Culture Facility), supplemented with 10% (vol/vol) FBS and 100 μg/ml penicillin/streptomycin (UCSF Cell Culture Facility). All cells were grown at 37°C and 5% CO2 in a humidified incubator. The plasmids encoding “mCherry-Vimentin-C-18” and “tdTomato-Clathrin-15” (the last number indicating the linker length between the fluorescent protein and the target protein) were purchased from the Michael Davidson Fluorescent Protein Collection at the UCSF Nikon Imaging Center. We transfected BSC-1 cells grown on an 8-well glass bottom chamber (Thermo Fisher Scientific) using FuGene HD (Promega). For better cell attachment, 8-well chamber was coated with fibronectin solution (Sigma-Aldrich, F0895-1MG) for 45 min before seeding cells. Cells were seeded at the density of 2000~4000 per well one day before transfection. Total plasmid amount of 200 ng per well with the clathrin to vimentin ratio in 1:4 was used to achieve optimal two-color labeling. Forty-eight hours after transfection, cells were fixed with 4% paraformaldehyde for 15 mins, followed by three times of PBS washes.

In order to minimize the background fluorescence, our hyperspectral detection setup was built around a wide-field inverted fluorescence microscope (Nikon Ti-U) with an oil immersion objectives (Olympus 100x 1.4 Plan Apo). Excitation light beans (488 nm and 561 nm, Coherent OBIS) were firstly combined and expanded, then pass through a home-built total internal reflection fluorescence (TIRF) illuminator before entering the microscope. The incident angle of the excitation light can be adjusted to be just smaller than the critical angle. Fluorescence was filtered using a quad-band dichroic mirror (z405/488/561/640rpc, Chroma) before exiting the microscope body. A pair of lenses (Fig. 1a, L1: f = 150 mm, L2: f = 100 mm) relayed the intermediate image plane to the camera, with the optical elements for spatial-spectral dual view imaging placed in between. Images were recorded by a sCMOS camera (ORCA Flash 4.0 sCMOS, Hamamatsu). The final pixel size at the image plane is 85 nm. During imaging, camera frame rate was set to be 5 Hz.

## 3. Results

### 4.1 Evaluation using simulated two-color data

We first quantified the performance of the reconstruction algorithm by analyzing a simulated data set (see Section 2.4) (Fig. 2). Experimentally acquired images of microtubule (Fig. 2a) and mitochondria outer membrane protein Tom20 (Fig. 2b) were assumed to be labeled by EGFP and EYFP. Conventionally, EGFP and EYFP are imaged in the same green channel, because their emission peaks are separated by only 20 nm (Fig. 2c). Here we used this probe pair to demonstrate the reconstruction accuracy and spectral resolution of our computational method. For this purpose, simulated images (Figs. 2d, e) with various SNR and SBR were generated. We set the average signal level to be between 100 and 1500 photons and SBR to be between 0.1 and 1. We note that in wide field imaging, the only significant noise source is Poisson noise. Other noises such as camera readout noise are negligible at realistic levels of signal. As the result, the average SNR of the simulated dataset is the square root of average signal level. We purposely chose relatively low signal levels to examine the effect of noise on the reconstruction algorithm.

**Fig. 2.**
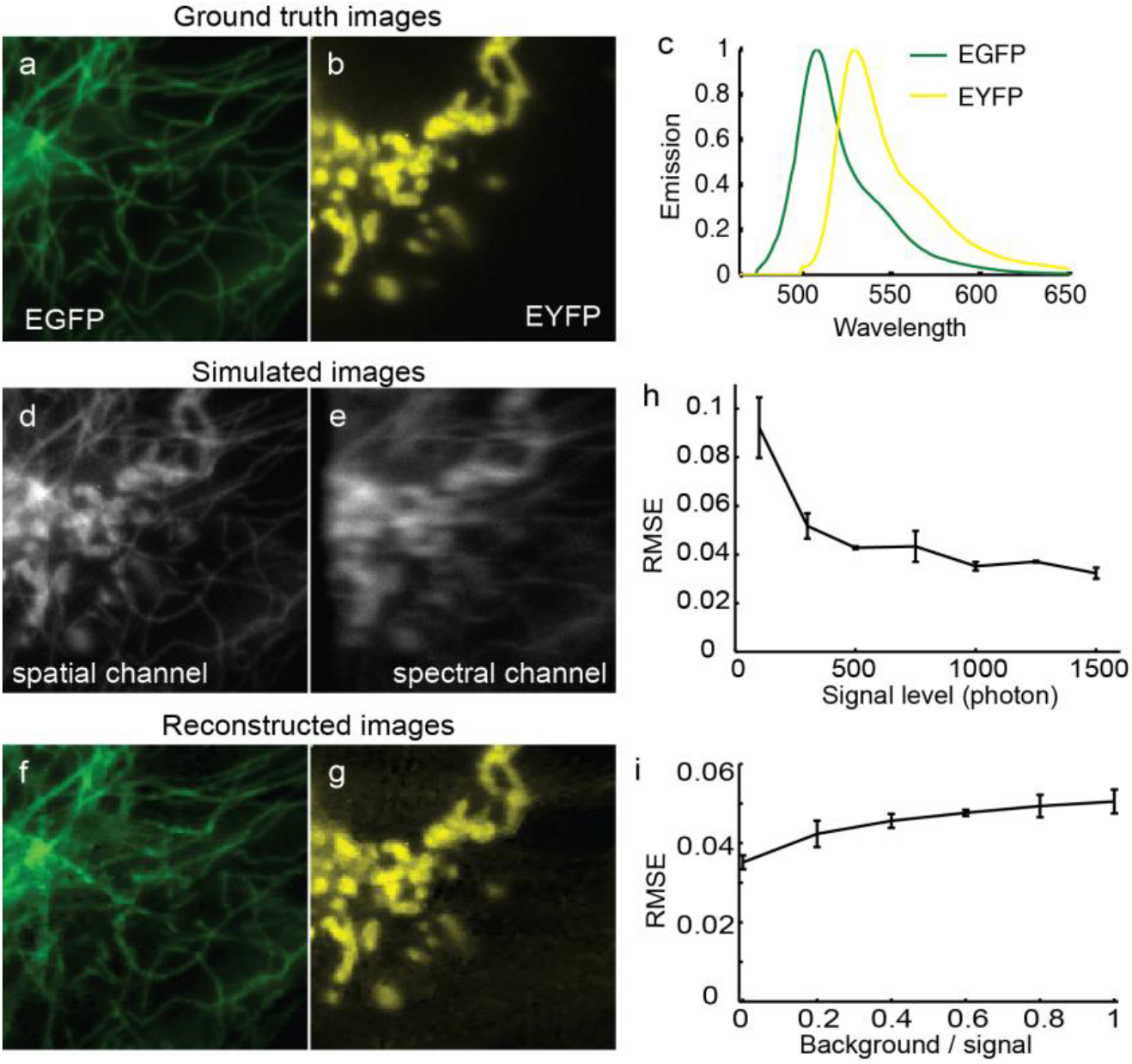
Simulated results of two-color imaging using space-constrained computational hyperspectral imaging. (a, b) “Ground truth” microtubule and mitochondria images. Image size: 128 x 128 pixels. (c) The normalized emission spectrum of EGFP and EYFP, showing the emission peak separation is about 20 nm. (d, e) Simulated spatial (d) and spectral (e) images by assuming that microtubule and mitochondria are label by EGFP and EYFP respectively. (f,g) Representative reconstructed images of the two color channels showing clean signal separation. Average signal level and background were set to 1000 and 300 respectively. (h) The dependence of reconstruction root mean square error (RMSE) of our method on average signal level when there is no background. (i) The dependence of reconstruction RMSE on different amount of homogeneous background. The average signal was 1000.

We compared the reconstructed results (Figs. 2f, g) with the ground truth images (Figs. 2a, b) and quantified the accuracy by the root mean squared error (RMSE) between them. In order to measure RMSE independent of the absolute signal level, we used the relative RMSE, which was calculated after the ground truth and reconstructed images are normalized to its maxima. Fig. 2h shows that our method had a relative RMSE smaller than 0.1 even when signal level in extremely low. For signal level higher than 500, our method steadily presented a RMSE of ~0.04. Besides, our method had robust performance (0.03 ~ 0.05 relative RMSE) under a wide range of background signal level. With this performance, our method was able to accurately separate EGFP and EYFP with emission peaks separated by ~20 nm. We noted that the main contributor to this performance is the space-constraint from the spatial image. Without space-constraint, the algorithm failed to correctly assign signals between channels.

### 4.2 Analysis of two-color experimental data

We then validated our method in analyzing real experimental images. Here we demonstrated two-color imaging by tdTomato-clathrin and mCherry-vimentin. This sample was chose for two reasons. First, because the true underlying two channels were unknown, two distinct structures were chosen to facilitate visual evaluation of reconstruction performance. As shown in Fig. 3a and 3b, which are the acquired spatial and spectral images of the sample, the bright dots corresponds to clathrin and the filament structure is vimentin. By comparing the reconstruct results with the structural prior knowledge, we could better estimate the performance of our method in analyzing real experimental data. Second, tdTomato and mCherry were used because their emission peaks are separated by 19 nm (Fig. 3c). We note that even though the emission peak separation is comparable to that between EGFP and EYFP (Fig. 2c), it is more challenging to separate tdTomato and mCherry because their emission spectrum is much wider than EGFP and EYFP.

**Fig. 3:**
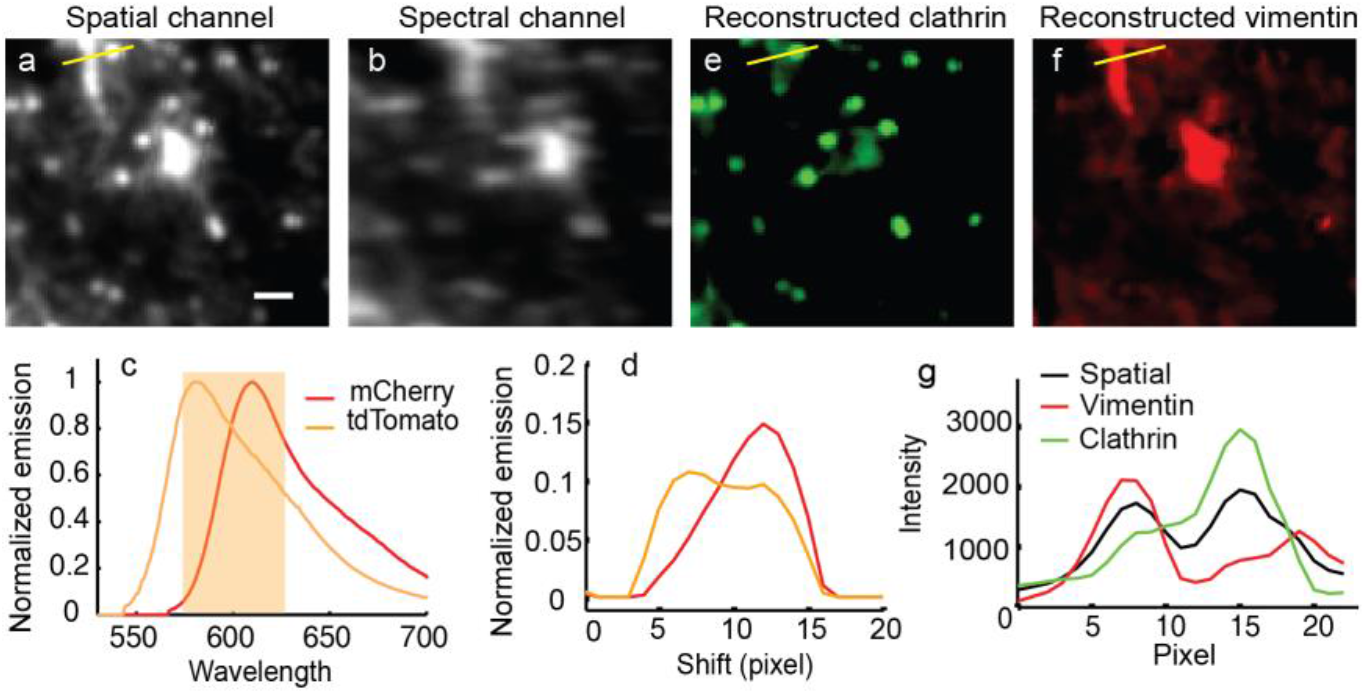
Experimental results of two-color imaging using space-constrained computational hyperspectral imaging. Images shown here are BSC-1 cells having clathrin and vimentin labeled with tdTomato and mCherry respectively. (a, b) Experimental images from spatial and spectral image paths. Scale bar: 1 μm. (c) Emission spectrum of mCherry and tdTomato, the orange shaded area shows the region of detection filter used in our system. (d) Calibrated emission from single-color imaging experiments. (e, f) Reconstructed images for tdTomato-clathrin (e) and mCherry-vimentin (f) respectively. (g) The intensity profile along the yellow line in (a,d,e), showing the algorithm successfully separates clathrin and vimentin structures.

Using dual-view registration function (obtained as described in section 2.4) and dispersed convolution kernel functions (Fig. 3d) calibrated from single-color labeled sample, we successfully reconstructed clathrin (Fig. 3e) and vimentin structure (Fig. 3f) with low cross talk. Vimentin filaments were well resolved despite that their intensity varied a lot in the field-of-view (FOV). Most of the clathrin dots were also well resolved except the cross-talk at upper left corner and center of FOV where vimentin signal was strong. The intensity profile plotted in Fig. 3g quantitatively shows that the resolution was not sacrificed during reconstruction, evidenced by the peak width is very similar between two reconstructed clathrin and vimentin channels and the spatial image. We noted that even though the emission of tdTomato and mCherry can potentially be separated by dichroic mirror-based ratio-metric imaging, it usually requires change or optimization of dichroic mirror according to different probes. Our method, on the other hand, has the advantage of totally free wavelength choice.

### 4.3 Evaluation using simulated three-color data

As mentioned in section 2.1, when the sample is multi-color labeled, multiple measurements of the sample using random illumination patterns generated by DMD can be used to avoid under-sampling of the 3D data. We illustrated the proposed setup diagram (Fig. 4a), in which DMD was placed in the excitation path, at the conjugate plane of the sample plane. Placing DMD in the detection path would result in significant loss of emitted photons. As a proof-of-principle, we used simulation to demonstrate the effectiveness of our computational hyperspectral imaging method in three-color imaging. Based on the simulation of Section 3.1, we added another clathrin-RFP channel. Random illumination patterns were simulated by generating random binary patterns of the same size with the “ground truth” images (Fig. 4b), and then these patterns were multiplied to ground truth and measurement matrix before generating simulated spatial and spectral images (Fig. 4a). Fig. 4c displays the clean separation of all the three channels, with the cross-talk slightly higher than two-color imaging. The reconstruction performance depends on the number of illumination patters. We found that for three-color imaging, relative RMSE decreased from ~0.1 to ~0.04 with the increasing of the number of illumination patterns from 1 to 10 (Fig. 4d). This result means that more measurements ensures a higher reconstruction accuracy. However, more measurements also lowers the temporal resolution of computational hyperspectral imaging and makes this method less advantageous compared to scanning hyperspectral imaging. This tradeoff indicates we need to choose a relative optimal number of illumination patterns. For three-color imaging, a pattern number of five is a good tradeoff between reconstruction accuracy and imaging speed (Fig. 4d).

**Fig. 4:**
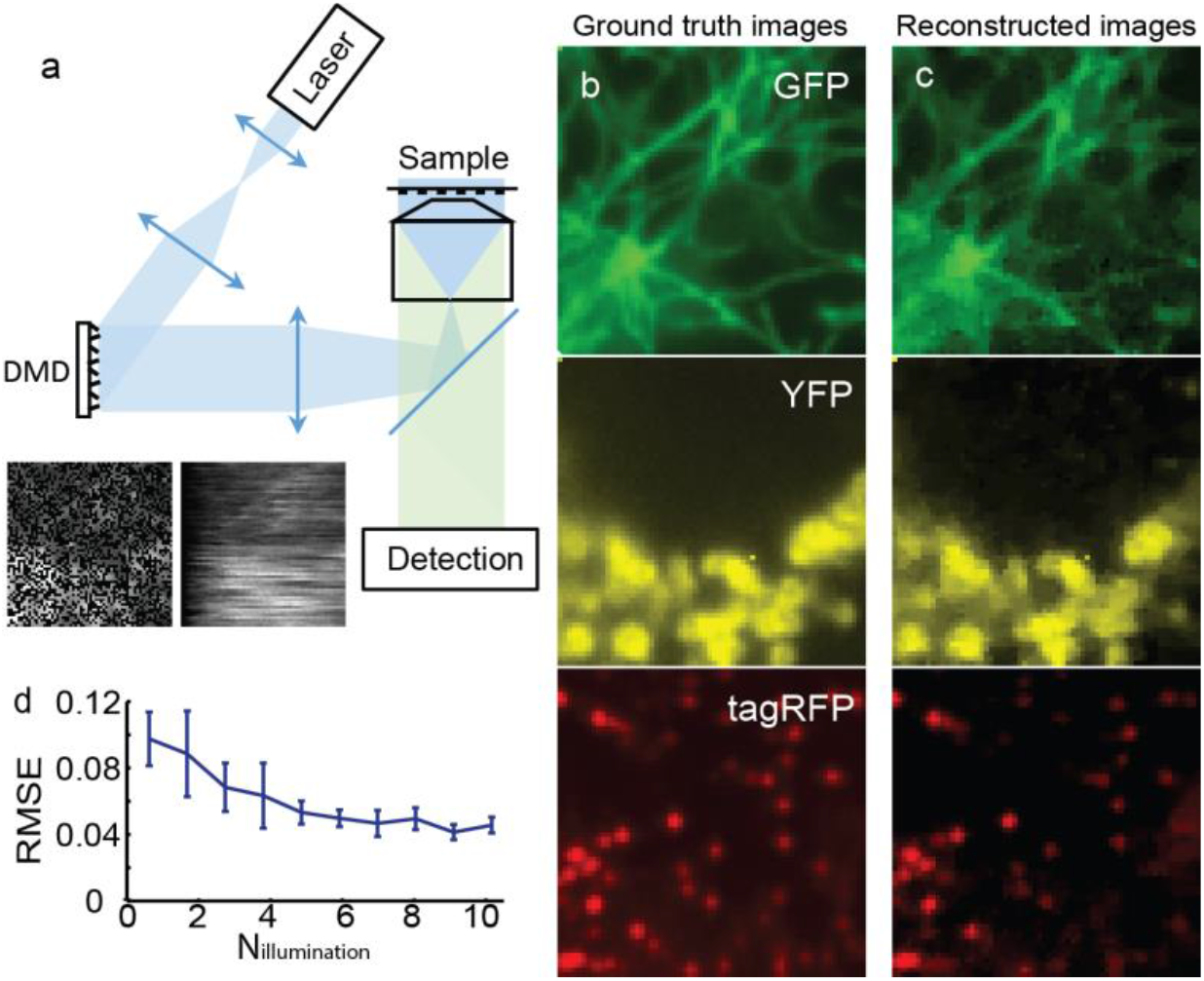
Multi-color computational hyperspectral imaging scheme. (a) Illustration of excitation scheme, in which DMD is placed at the conjugate plane of the sample to generate random illumination.

Representative simulated spatial (left) and spectral (right) images after adding a random illumination pattern to the ground truth were shown. (b) “Ground truth” microtubule, mitochondria and clathrin images. Image size: 64 x 64 pixels. (c) Representative reconstructed images of the three color channels. Average signal level and background was set to 1000 and 300 respectively. The number of illumination patterns is 5. (d) Dependence of reconstruction RMSE on the number of illumination patterns.

## 4. Conclusion

We presented a computational hyperspectral imaging method for wide-field multi-color fluorescence imaging. The method is based on simultaneous spatial and spectral data acquisition, in which the spatial information acted as a constraint in the image reconstruction to achieve superior accuracy. With both simulation and experimental data, we have shown that our method has a high reconstruction accuracy in separating probes with emission peak separation of 20 nm for two-color imaging. As a proof-of-principle, we have also shown with simulation that our method has the multi-color imaging capability given multiple measurements from patterned illuminations. Our method provides completely free wavelength choice for multi-color imaging. It is particularly advantageous when combined with conventional filter-based sequential multi-color imaging to further expand the probe choices of fluorescent imaging.

The same method can also be combined with other microscope beside TIRF, for example spinning disk confocal and light sheet microscopy. We note that the main limitation of our method is that the application is confined to imaging distinct structures. Relatively homogeneous distributed protein over the FOV, for example diffuse membrane protein, would result in homogeneous spectral image, and the algorithm would fail to extract information from it.

## Acknowledgement

We thank Grace Guo from Laural Waller’s lab for the fruitful discussion regarding the reconstruction algorithm.

